# A multi-platform reference for somatic structural variation detection

**DOI:** 10.1101/2020.10.15.340497

**Authors:** Jose Espejo Valle-Inclan, Nicolle J.M. Besselink, Ewart de Bruijn, Daniel L. Cameron, Jana Ebler, Joachim Kutzera, Stef van Lieshout, Tobias Marschall, Marcel Nelen, Andy Wing Chun Pang, Peter Priestley, Ivo Renkens, Margaretha G.M. Roemer, Markus J. van Roosmalen, Aaron M. Wenger, Bauke Ylstra, Remond J.A. Fijneman, Wigard P. Kloosterman, Edwin Cuppen

**Affiliations:** Center for Molecular Medicine and Oncode Institute, UMC Utrecht, The Netherlands; Hartwig Medical Foundation, Amsterdam, The Netherlands; Bioinformatics Division, Walter and Eliza Hall Institute of Medical Research, Melbourne, Australia; Institute for Medical Biometry and Bioinformatics, Medical Faculty, Heinrich Heine University Düsseldorf, Germany; Department of Human Genetics, Radboud UMC, Nijmegen, The Netherlands; Bionano Genomics, San Diego, California, USA; Department of Pathology, Amsterdam UMC, Vrije Universiteit Amsterdam, Cancer Center Amsterdam, The Netherlands; Pacific Biosciences, Menlo Park, California, USA; Department of Pathology, Netherlands Cancer Institute, Amsterdam, The Netherlands

**Author notes:** **Correspondence:** /.

## Abstract

Accurate detection of somatic structural variation (SV) in cancer genomes remains a challenging problem. This is in part due to the lack of high-quality gold standard datasets that enable the benchmarking of experimental approaches and bioinformatic analysis pipelines for comprehensive somatic SV detection. Here, we approached this challenge by genome-wide somatic SV analysis of the paired melanoma and normal lymphoblastoid COLO829 cell lines using four different technologies: Illumina HiSeq, Oxford Nanopore, Pacific Biosciences and 10x Genomics. Based on the evidence from multiple technologies combined with extensive experimental validation, including Bionano optical mapping data and targeted detection of candidate breakpoint junctions, we compiled a comprehensive set of true somatic SVs, comprising all SV types. We demonstrate the utility of this resource by determining the SV detection performance of each technology as a function of tumor purity and sequence depth, highlighting the importance of assessing these parameters in cancer genomics projects and data analysis tool evaluation. The reference truth somatic SV dataset as well as the underlying raw multi-platform sequencing data are freely available and are an important resource for community somatic benchmarking efforts.

## Introduction

Structural genomic variations (SVs) form a major class of somatic genetic variation in cancer genomes (Li et al., 2020; Yang et al., 2014), involving dozens to thousands of somatic SVs with varying size distribution and patterns (Li et al., 2020). While some SVs represent simple deletions, others tend to be complex, involving multiple breakpoints across a relatively short genomic interval. For example, chromothripsis is a form of complex SVs frequently observed in cancer genomes (Cortés-Ciriano et al., 2020; Kloosterman et al., 2014), resulting from aberrant chromosome segregation or telomere dysfunction (Maciejowski et al., 2015; Zhang et al., 2015). Other types of complex SVs involve oncogene amplifications arising from breakage-fusion-bridge cycles (Bignell et al., 2007; Li et al., 2020; Nattestad et al., 2018). SVs in cancer genomes may promote cancer development through a variety of mechanisms, such as oncogene activation through gene-fusions, disruption of tumor suppressor genes or by affecting gene regulation (Mitelman et al., 2007; Spielmann et al., 2018). Oncogenic fusion genes resulting from somatic SVs form important targets for cancer drugs, and somatic SVs may form neo-antigenic targets for immunotherapies (Mansfield et al., 2019), demonstrating the relevance of accurate somatic SV detection for personalized cancer treatment (Mertens et al., 2015; Mitelman et al., 2007).

While classical karyotyping and FISH analyses have been instrumental in systematic copy number analyses in tumor samples (Mertens et al., 2015; Mitelman et al., 2007), these technologies provide limited resolution or do not allow for comprehensive genome-wide analysis and are thus unable to resolve the complete spectrum of SV events. Most of our knowledge on genome-wide high-resolution SVs in cancer genomes stems from the analysis of short-read whole genome sequencing, which is currently the only scalable and cost-efficient technology for high-resolution genome-wide cancer genome analysis (Li et al., 2020; Macintyre et al., 2016). Although short reads are effective for detection of simple SV breakpoints in non-repetitive regions of the genome, the interrogation of complexly rearranged regions or the detection of SV breakpoints in low complexity genomic regions may require other sequencing techniques or targeted approaches (de Vree et al., 2014). For example, long-insert mate-pair sequencing has proven a valuable strategy for genome-wide mapping of somatic SVs (Hillmer et al., 2011; Kloosterman et al., 2011) and single-cell template strand sequencing enables the detection of copy number variants and copy neutral structural variants (Sanders et al., 2020). Furthermore, long-read sequencing methods, such as Pacific Biosciences and Oxford Nanopore and synthetic long-read approaches, such as linked-read technology by 10x genomics, provide a promising alternative for the detection of SVs. Initial studies have shown that long-read single-molecule sequencing can greatly improve detection of germline SVs (Chaisson et al., 2014, 2019; Cretu Stancu et al., 2017; Huddleston et al., 2017). Similarly, recent work has demonstrated the advantage of long-range sequence information for identification of SVs in cancer genomes, such as cancer gene amplifications and gene fusion events (Greer et al., 2017; Gupta et al., 2015; Nattestad et al., 2018; Zheng et al., 2016).

A major limitation of studies on cancer SVs is the lack of a comprehensive ground truth genome-wide somatic SV datasets including all types and sizes of somatic structural aberrations. Such truth sets can form a resource for benchmarking sequencing and analysis approaches as well as for evaluating detection problems related to intratumor heterogeneity and tumor purity. Gold reference truth sets have been established for germline SVs (Chaisson et al., 2019; Zook et al., 2020) or somatic single nucleotide variants (SNVs) (Arora et al., 2019). However, attempts at benchmarking somatic SVs have only been performed by using *in silico* simulated data (Gong et al., 2020; Lee et al., 2018), or mouse data (Sarwal et al., 2020).

We addressed this caveat by generating a multiplatform short-read, long-read and linked-read sequencing and optical mapping dataset for the COLO829 melanoma cell line and the paired COLO829BL lymphoblastoid reference cell line. These cell lines have been used before to establish somatic SNV and copy number alteration (CNA) reference sets (Arora et al., 2019; Craig et al., 2016; Pleasance et al., 2010). By cross-platform comparison and extensive validation we define a gold reference set of 68 somatic SVs in COLO829. We evaluated the completeness of this validated truth set and demonstrated its use to study the effect of tumor purity and sequencing coverage variation on the accuracy of somatic SV calling. We believe this somatic SV truth set to be of broad value for benchmarking and quality control of large-scale cancer genome sequencing studies, which are currently undertaken in research and the clinic.

## Results

### Multi-platform genome-wide analysis of the COLO829 tumor-normal melanoma cell line pair

In this study, we aimed at obtaining a comprehensive view on the genome structure of the COLO829 cancer cell line and identify a high-quality set of somatic structural variations, for use as a reference dataset. We cultured COLO829 and the corresponding lymphoblastoid cell line (COLO829BL) according to standard conditions (**Materials and Methods**). A large batch of cells expanded from one original vial directly obtained from the ATCC cell line repository was used for DNA isolation and subsequent genomic analysis using five different technology platforms: Illumina HiSeq Xten (ILL), Oxford Nanopore Technologies (ONT), Pacific Biosciences (PB), 10x genomics (sequenced on Illumina NovaSeq; 10X), and Bionano Genomics Saphyr optical mapping (BNG) (**Materials and Methods**).

The sequencing and optical mapping data were analyzed with respect to the reference human genome (GRCh37) using alignment methods specific for each technology (**Materials and Methods**). From the combined short and long read sequencing data of the COLO829 sample we obtained a total average base coverage of 235X, while the BNG data generated an additional physical coverage of 218X. For the COLO829BL control cell line a combined average base coverage of 155X and a BNG physical coverage of 220X was reached (**Figure 1A, Supplementary Table 1**). Average physical molecule lengths were 534 bp for ILL paired-end inserts, 11 kb for ONT, 19 kb for PB and 98 kbp for BNG optical maps (**Figure 1B, Supplementary Table 1**).

**Figure 1.**
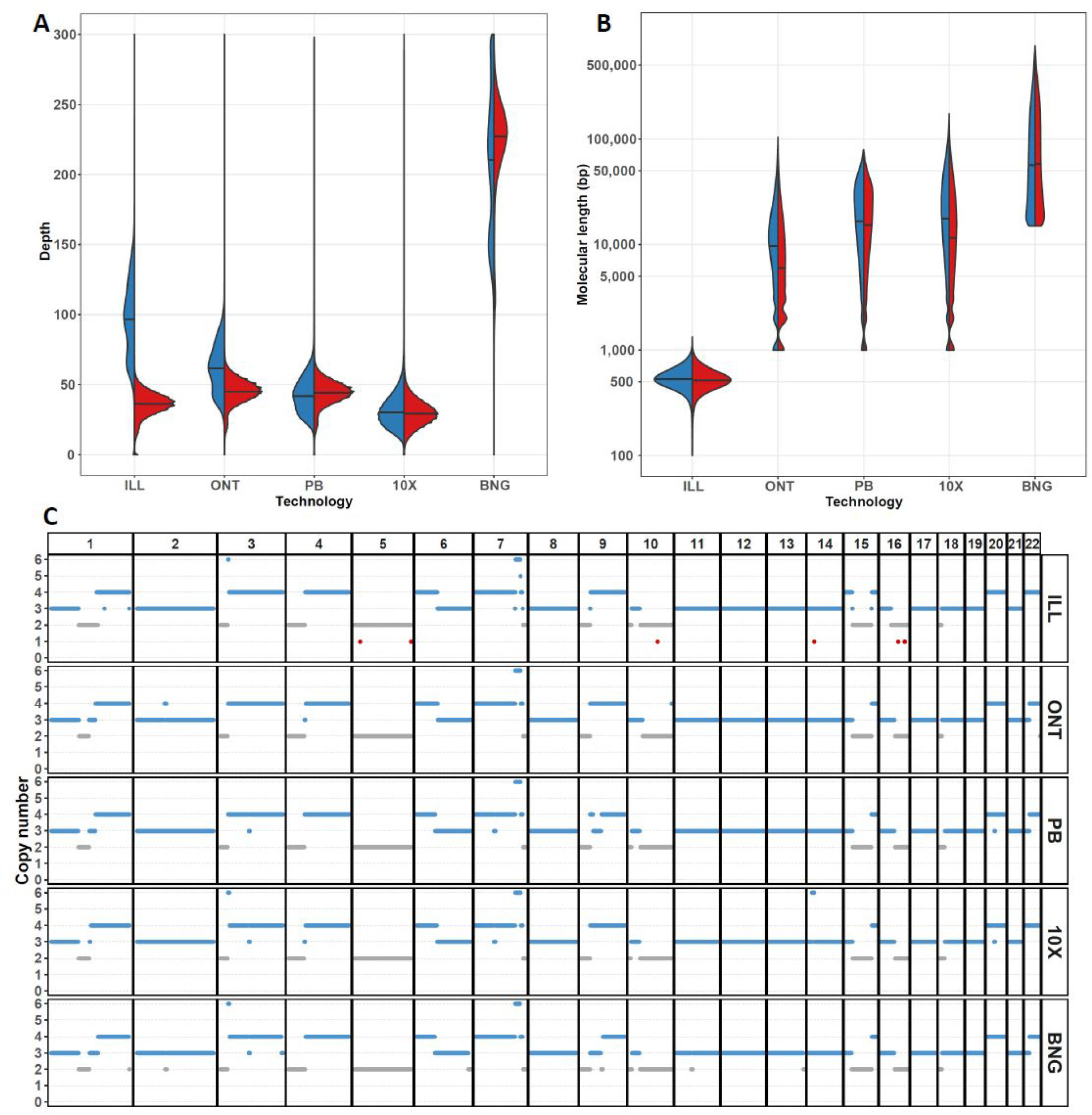
Overview of the COLO829 multi-technology genomic dataset. Sequencing depth **(A)** and log-scaled molecular analysis length **(B)** distributions per technology dataset for COLO829 (blue) and COLO829BL (red). Means are indicated by horizontal black lines. **(C)** Copy number profile of COLO829 calculated independently for each of the datasets.

To assess the consistency of each of the technologies with respect to representation of the sequence content of the COLO829 cancer cell line, we determined the presence of copy number alterations. This revealed a highly similar copy number profile for each of the technologies (**Figure 1C**), with a correlation of copy number calls in the different datasets of 0.87-0.96 (**Supplementary Figure 1A**). Furthermore, we compared our copy number calls with those generated in previous bulk (Arora et al., 2019) and single cell (Velazquez-Villarreal et al., 2020) sequencing of COLO829. The overall CNA landscape of the bulk sequencing and the dominant cluster from single cell sequencing is very similar to the one we obtained (**Supplementary Figure 1B**), with a correlation of 0.99 (bulk) and 0.97 (single cell group A) (**Supplementary Figure 1C**). However, the previously described subclonal single cell clusters (B-D) possess some distinct copy number aberrations that are not observed in our bulk sequencing datasets (i.e. extra copy of chromosome 8 in group D or lack of gain in short arm of chromosome 1), in line with the proposed continuous genomic evolution of cell lines and subculture-specific nature of these events. Finally, classical FISH analysis for six genomic locations of the culture used in our study confirmed the sequencing derived chromosomal copy number states (**Supplementary Figure 3D**).

### Generation of a somatic structural variation consensus truth set

To build an accurate and comprehensive somatic SV truth set, we used a combinatorial analysis approach involving the four sequencing platforms (ILL, ONT, PB and 10X). Somatic SVs were obtained using state-of-the art SV calling approaches defined for each of the sequencing datasets **(Materials and Methods, Figure 2A**). SV calling parameters were not necessarily optimized for highest precision, but to high sensitivity to not miss out on any real event. As a result, individual candidate call sets for each technology resulted in highly variable lists of predicted somatic SVs, ranging from 92 breakpoint calls in ILL up to 6,412 for ONT, adding up to a total of 8,831 merged candidate somatic SV calls **(Figure 2A)**. Only 18 of those somatic SV calls were found by all four sequencing approaches and 125 SV calls were supported by at least two call sets (**Supplementary Figure 2A**).

**Figure 2.**
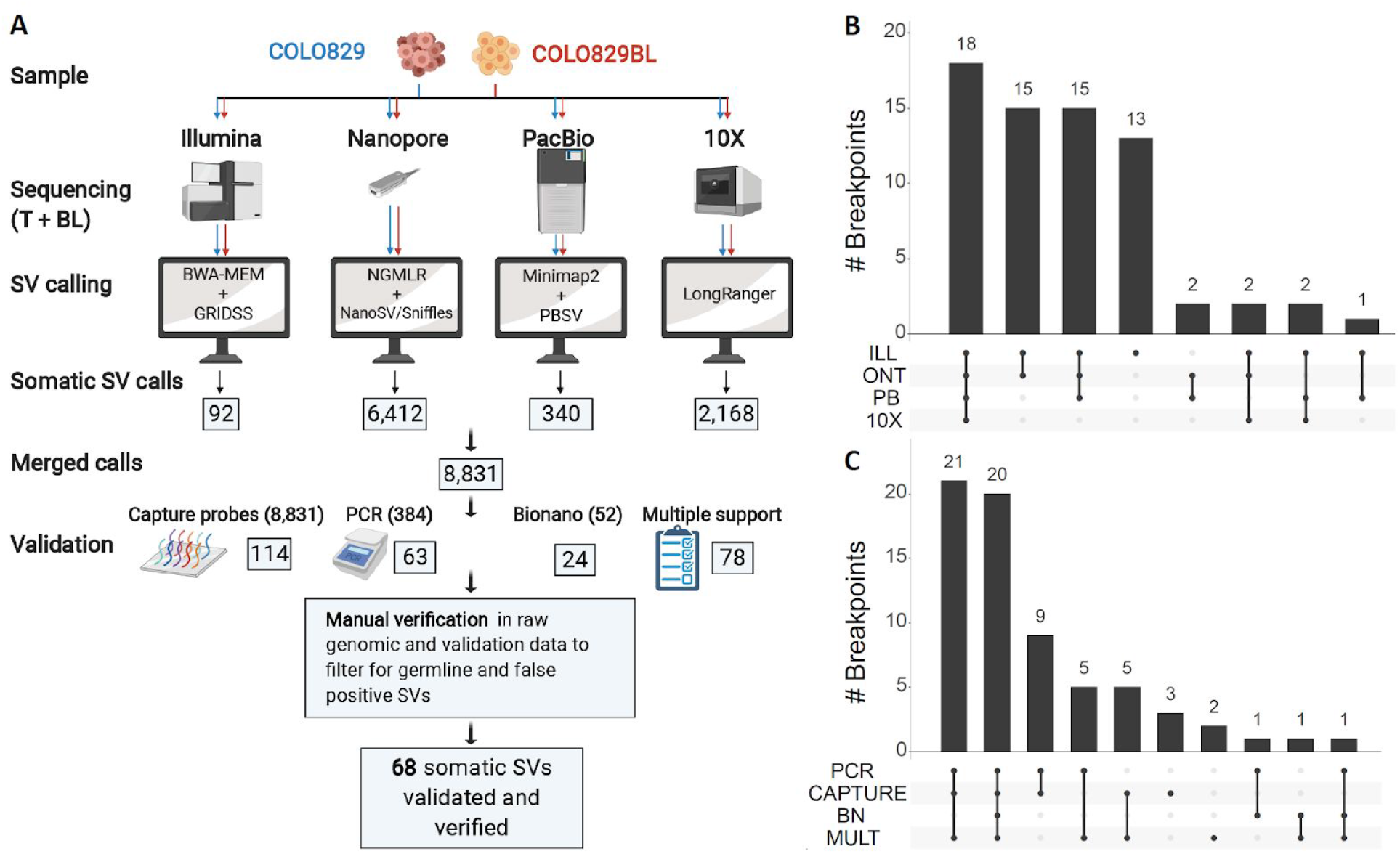
Generation of a validated somatic SV truth set. **(A)** State-of-the-art somatic SV calling pipelines were used independently for each technology dataset. The number of somatic SV candidates identified are indicated in boxes. Overlapping variant calls obtained by the different platforms were merged and independently validated using a combination of targeted enrichment with hybrid capture probes followed by next-gen sequencing, PCR and Bionano genomics. Validated somatic SV candidates and calls supported by more than one dataset were manually curated, leaving a total of 68 somatic SVs in the truth set. Intersections between the 68 somatic SVs in the truth set and the original SV call sets **(B)** and the validation results **(C)** are shown. ILL = Illumina HiseqX, ONT = Oxford Nanopore, PB = PacBio, 10x = 10x Genomics, BN = Bionano, MULT = support by multiple sequencing platforms.

To make an initial assessment of accuracy, we selected 88 high-confidence SV candidates for PCR validation based on visual inspection of the mapped reads using IGV. In addition, we randomly selected 296 additional SV candidates for PCR validation. Based on short and long read sequencing of the PCR products, 63 of these breakpoints were labelled as PCR validated **(Supplementary Figure 2B)**. Moreover, we decided to perform a separate validation of all 8,831 somatic SV calls from the union of the four SV callsets, using a capture-based enrichment method using multiple probes flanking and overlapping each candidate break-junction (**Materials and Methods**). Based on the short read sequencing of the enriched products, 114 breakpoints were labelled as capture validated **(Supplementary Figure 2B)**. Lastly, we used the 52 BNG somatic SV calls as an additional layer of validation. In total, 137 SV calls were validated by at least one of the methods aforementioned. Additionally, 78 SV calls were not validated but still supported by more than one technology. (**Figure 2A, Supplementary Figure 2C**).

Next, we manually curated these 215 SV calls that were either validated or supported by multiple technologies. Based on visual inspection of the genomic alignment data from each of the sequencing sets and the validation experiment results, we assessed each SV call individually. We found that 14 calls were actually duplicate calls of the same event (but annotated slightly different by different data analysis pipelines), 48 were real events but also had evidence in the germline control, and another 98 were considered false positive as the supporting or reference data was very noisy at the given genomic location (also in the independent validation data) and may thus reflect the impact of low confidence regions in the reference genome for which unambiguous mapping of sequencing reads is complicated due to simple sequence or repeat content. Taken together, we conclude that 68 of the SV candidates are real somatic events and thus considered our truth set (**Figure 2A, Supplementary Figure 2C, Supplementary Table 2 with all validations and raw calls**). To verify the efficacy of our manual curation approach, we randomly selected 179 SV calls that were supported by a single technology and not validated, and therefore left out from the candidate SV curation pipeline, and also evaluated them manually. All these SV calls were either germline events (63, 35%) or false positive due to noisy mapping data (116, 65%) (**Supplementary Figure 2D**).

Of the compiled set of 68 validated somatic SVs in COLO829, 55 (81%) were present in more than two original call sets, including the 18 SVs detected by all technologies (**Figure 2B**). Moreover, most of the SVs were validated at least by capture-based enrichment and by PCR (50, 74%). Additionally, 8 somatic SVs were validated by capture-based enrichment but not by PCR and vice versa, 7 somatic SVs were validated by PCR but not by capture-based enrichment. Of the remaining 3 SVs, one was validated by BNG and 2 were not validated by any targeted assay but are supported by multiple technologies and manually verified by inspection of raw sequencing data from both tumor and normal samples (**Figure 2C**). The resulting somatic SV truth set is presented in **Supplementary Table 3** and freely available as VCF.

### Characterization of the COLO829 somatic SV truth set

The somatic SV truth set consists of 38 deletions, 3 insertions, 7 duplications, 7 inversions and 13 translocations (**Figure 3A**). Most of the deletions (24, 61%) are larger than 10kbp, and 7 are smaller than 100bp. There are also three duplications and three inversions larger than 10kbp. Two tumor driver genes are affected by somatic SVs in COLO829 (**Supplementary Table 3**). First, there are two large heterozygous deletions (72 kb and 141 kb) in FHIT, located in the fragile site FRA3B on chromosome 3, which is commonly affected by somatic SVs (Li et al., 2020). Second, there is a homozygous 12 kbp deletion affecting PTEN on chromosome 10.

**Figure 3.**
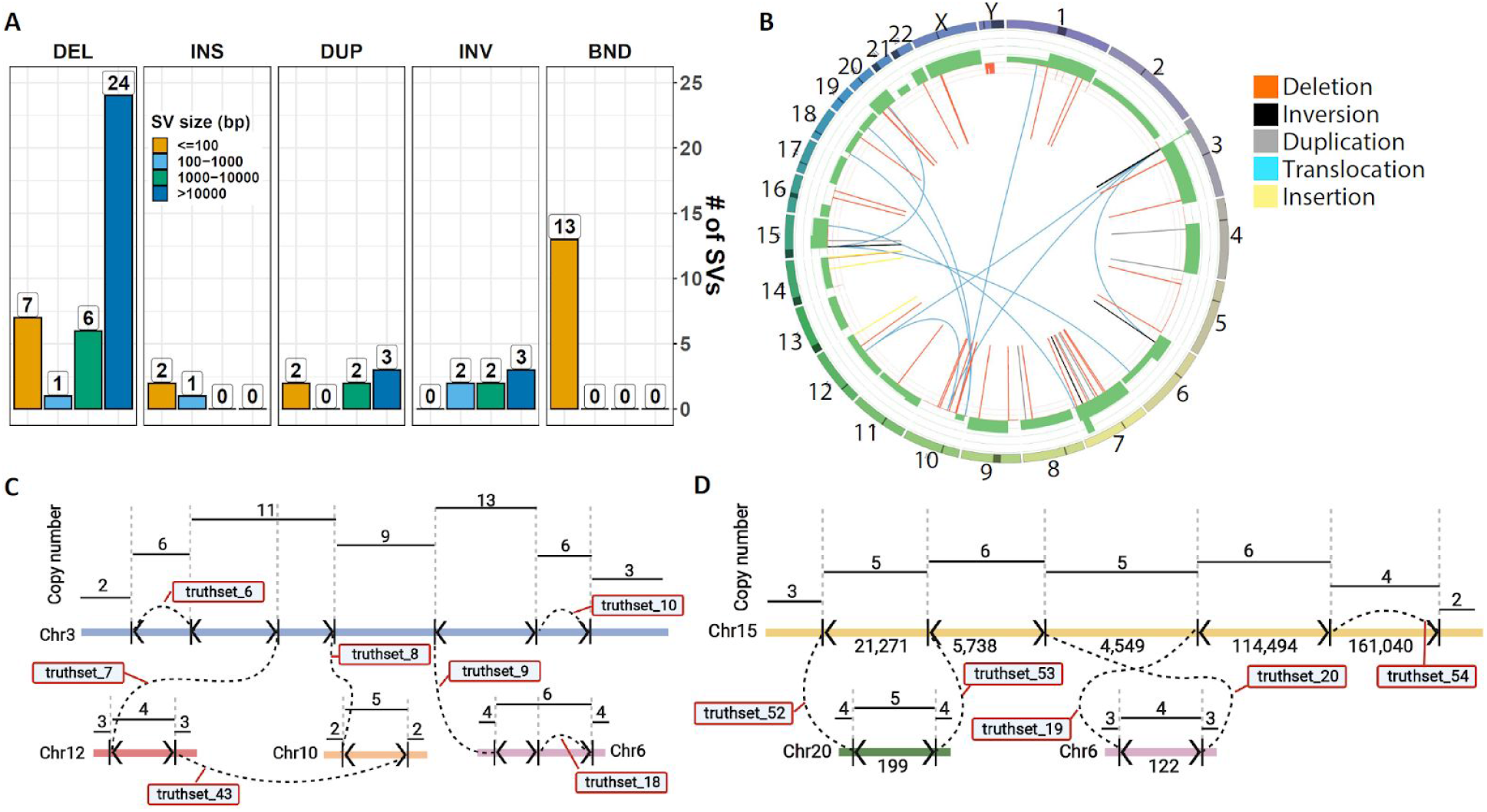
Characterization of the somatic SV truth set: **(A)** Distribution of different types of SVs in the COLO829 truth set, divided in size bins. Translocations (BND) are assigned a size of 0 bp. **(B)** Correlation between CNAs and somatic SVs in the COLO829 truth set. The circos plot shows copy number gains (green) and losses (red) and somatic SVs. Each copy number change is expected to be flanked by an SV event. Two complex breakage-fusion-bridge events are present in COLO829. The first one **(C)** occurs in chromosome 3 (blue), with templated insertions from chromosomes 6 (pink), 10 (green) and 12 (red) (see also Supplementary Movie for an animation of the proposed mechanism shaping this event). The second one **(D)** occurs in chromosome 15, with templated insertions from chromosomes 6 (pink) and 20 (orange). Breakpoints are indicated by vertical lines with arrowheads showing breakpoint orientations. Dashed lines indicate junctions between two breakpoints. Break-junctions are labelled with truth set SV ID number (Supplementary Table 3).

Frequently, SVs do not occur as simple isolated events but are part of a complex landscape induced in a single event like for example chromothripsis or due to a cascade of events over time like breakage-fusion-bridge cycles. There are also 2 clusters of complex chained somatic SVs that affect 3 or more chromosomes and involve more than 5 breakpoint junctions. Both of them resemble breakage-fusion-bridge events, since they are flanked by foldback inversions and show oscillating copy number profiles (Li et al., 2020). One of them occurred in chromosome 3 and involves four foldback inversions, two of which have templated insertions from chromosomes 10 and 12 and chromosome 6, respectively (**Figure 3C**). The breakpoint and copy number profile of chromosome 3 can be fully explained by 4 cycles of breakage-fusion-bridge followed by chromatid duplication through a whole genome doubling event. Initiated by replication of unrepaired double-stranded break, the unstable chromosome 3 (due to the presence of two centromeres in a single chromatid) underwent a further 3 more rounds of BFB with a fragment of chromosome 6 inserted prior to the third doubling cycle, fragments of chromosomes 10 and 12 inserted immediately after the fourth doubling cycle, and a stable state achieved after the final breakage through repair to one of the centromeres (**Supplementary Movie**). The other breakage-fusion-bridge event occurred on chromosome 15 and includes templated insertions from chromosomes 6 and 20 (**Figure 3D**). The donor locations of these templated insertions are not affected by SV events.

To evaluate the completeness of the somatic SV truth set, we compared it with the somatic CNA calls, since each CNA should have SV breakpoints or telomeres at either end. We found 43 total CNA breakpoints that are not telomeric ends of chromosomes. Of these, 26 (60%) are concurrent with an SV breakpoint. We evaluated the rest of the CNAs in the raw genomic data (**Supplementary table 4**). Six more copy number breakpoints (14%) are present in the germline, flanking heterozygous germline CNA events that are homozygous in the tumor through a somatic loss of the other allele. The SV break-junctions of these CNAs are germline and therefore not part of the truth set. Finally, there are 11 somatic CNA breakpoints (26%) not concurrent with an SV breakpoint. Five of these missing CNA breakpoints are located in a centromeric region (chromosomes 1, 4, 6, 14 and 16) and are likely due to a missing somatic SV involving the centromere, which are typically hard to fully resolve due to their repetitive nature. For another 2 missing CNA breakpoints (chromosome 3 and chromosome 9) breakends can be found in the raw ILL dataset, meaning an SV breakpoint was found but the SV junction partner could not be unequivocally determined. GRIDSS2 annotation did reveal that the chromosome 3 single break does map to one of the centromeres. Four more missing CNA breakpoints flank two supposed deletions in chromosome 1, but no SV call in these locations can be found for either COLO829 or COLO829BL in any of the datasets. Manual inspection of the raw data for these CNAs (**Supplementary Figure 3A, 3B**) indicates that these CNAs may actually reflect heterozygous germline events followed by LOH as witnessed by the lower read coverage in the COLO829BL as compared to the flanking segments. Furthermore, one CNA involves a LINE-rich region while the other overlaps with a segmental duplication.

Next, we compared our somatic SV truth set to the somatic SV calls presented by Arora et al. They provide two different somatic SV callsets, one generated by the HiSeq platform with 77 somatic SV calls and the other by the NovaSeq platform with 75 somatic SV calls. Since these were provided based on GRCh38 genomic coordinates, we lifted our somatic SV coordinates over to GRCh38. We found that 58 (75.34%) and 59 (78.6%) of the somatic SV calls for the HiSeq and the NextSeq callsets, respectively, overlapped with our somatic SV truth set on both sides of the SV (**Supplementary Figure 3**). We manually inspected the 20 non-overlapping somatic SV calls from the Arora et al dataset in our raw ILL, ONT and PB data (**Supplementary Table 5**). In the long-read raw data (ONT and PB) only 3 out of the 20 have some support (maximum 3 reads). In the ILL raw data, 9 out of the 20 have limited evidence, with only one or a few supporting reads. Only 4 of these 9 SV calls passed bioinformatic calling criteria in our original ILL somatic SV calls, but none of these were called by any other technology or independently validated by more sensitive PCR or targeted capture and deep-sequencing. Therefore we consider these candidates as technology-specific noise and were discarded from our truth set, although we can formally not exclude that these are real variants that are present at very low frequency (<1% in the sample). Finally, 13 SVs are present in our truth set and not in the Arora et al. data set. All were detected by at least two different sequencing techniques and independently validated.

### Effect of tumor purity and sequencing depth on somatic SV calling

To demonstrate the utility of the COLO829 somatic SV truth set, we evaluated the effect of tumor purity, which is highly variable amongst clinical samples, on SV calling. We used the available raw datasets and simulated tumor purities of 75% (TP75), 50% (TP50), 25% (TP25), 20% (TP20), and 10% (TP10) by random *in silico* mixing of the genomic data from COLO829 and COLO829BL for ILL, ONT and PB, respectively. We performed SV calling independently on each of these mixed sets and on the original tumor file (100% purity, TP100) and the normal file (0% purity, TP0). We then calculated the recall (percentage of truth set found) and precision (percentage of calls that belong to truth set). With the standard settings used, somatic SV recall and precision were found to be highly dependent on tumor purity for all three technologies. At TP75 and TP100, recall is the highest, with >94% for ILL, >67% for ONT and >65% for PB. With TP50, the recall slightly decreases to 90%, 52% and 61% for ILL, ONT and PB, respectively. For purities lower than TP50, the recall decreases further to <76%, <22% and <48% for ILL, ONT and PB, respectively. Precision follows a similar trend in the case of ILL, with precisions >78% for purities larger than TP50, and a drop to 63% in TP25. In the case of ONT and PB, the higher number of false positives impact severly on the precision rates, potentially reflecting maturity level of platform-specific tools for somatic SV detection in tumor-normal paired samples, but also presenting opportunities for further analysis parameter and tool optimisation.

Sequencing depth is another essential parameter to consider in tumor sequencing projects as it represents a trade-off decision between variant detection sensitivity and costs. To investigate the effect of sequencing depth in combination with tumor purity in somatic SV detection, we took one of the triplicates from each of the simulated ILL tumor purities (98x coverage) and subsampled them to 50x, 30x, 10x, 5x and 1x depths. We again performed somatic SV calling using the same standard pipeline on each of these simulated sets and calculated recall and precision. We observed that for depths of 50x and 98x and tumor purities over 50% recall was over 82%. In the case of 98x, even at TP20 a recall of 71% could be obtained, whereas for 50x at TP25 the recall decreased to 42%. For 30x sequencing depth, at TP100 recall was 84%, but at TP50 there was a decrease to 54% and at TP25 further to 10%. For lower coverages, recall was low. Surprisingly, depths of 30x and 50x had a higher precision at all tumor purities than 98x, with precision around 95% over TP50, compared to approximately 70% for 98x. While this could in theory be explained by the presence of subclonal SVs that are not included in the reference truth set but become detectable at higher sequencing depth, this might also be caused by stochastic effects due to increased measurement noise at higher sequencing depth which increases the number of false positive and therefore reduces precision (although recall is not affected). Further optimization of analysis tools and settings and deeper sequencing may resolve these issues.

**Figure 4.**
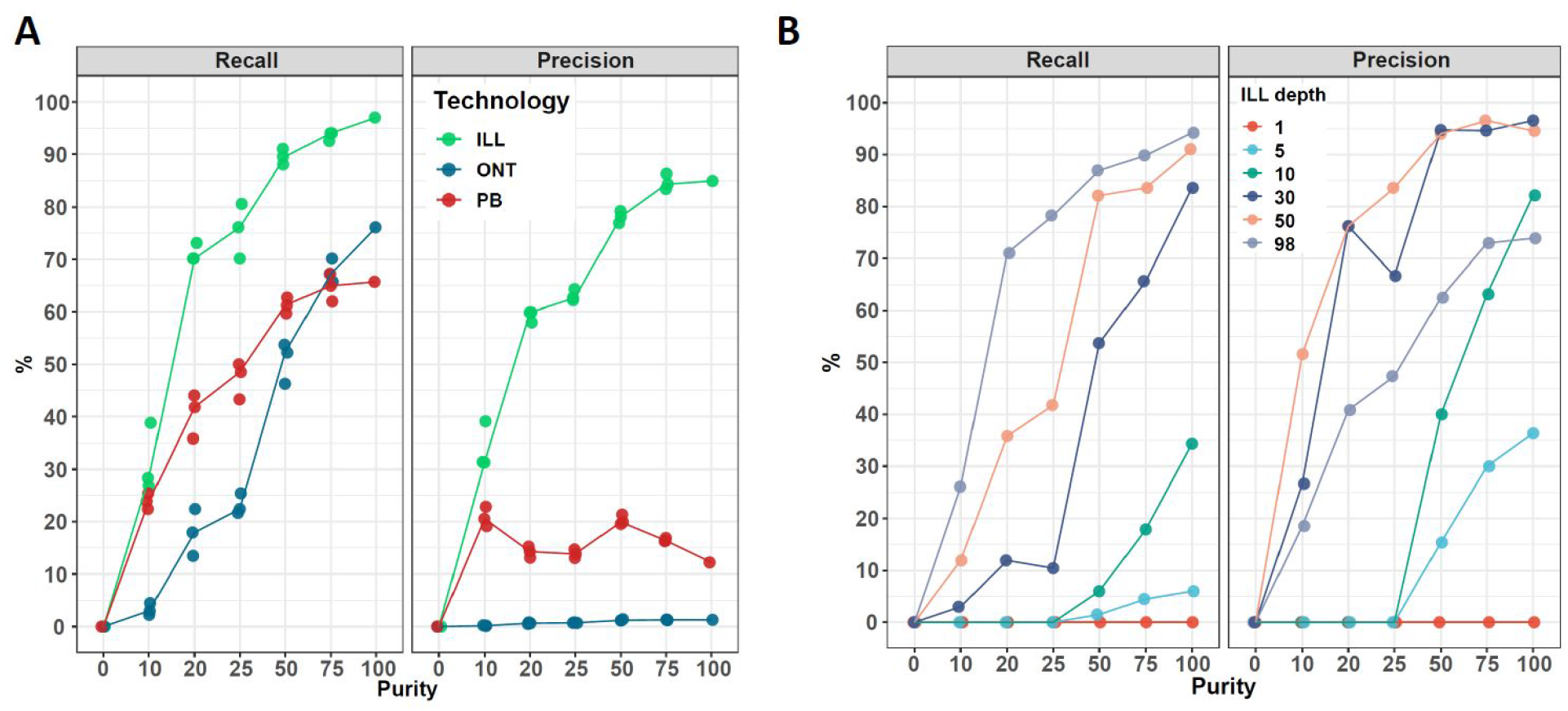
Recall and precision of somatic SV calling as function of tumour purity and sequencing depth effect. Different tumor purities (0, 10, 20, 25, 50, 75 and 100 %) were simulated by mixing data from COLO829 and COLO829BL for the ILL, ONT, and PB datasets. **(A)** Somatic SV calling was performed independently for each purity subset and recall (left) and precision (right) were calculated against the COLO829 somatic SV truth set. Lines represent the median of independent triplicate measurements. **(B)** For each tumor purity subset in the ILL dataset, different sequencing depths (1,5, 10, 30, 50 and 98x) were sampled. Somatic SV calling was performed independently for each sequencing depth and tumor purity subset and recall (left) and precision (right) were calculated against the COLO829 somatic SV truth set.

## Discussion

We produced a validated somatic SV truth set by building upon the strengths of different sequencing technologies. Bioinformatic integration of results and large-scale independent validation strategies turned out to be a powerful approach for reducing the large number of candidate events obtained. Manual curation and inspection of raw sequencing data was however essential to exclude sequencing, mapping artefacts and remaining germline events. These somatic false positives are thus germline false negatives and were likely included in the initial somatic SV calls due to the lower sequencing analysis depths for the control sample as compared to the tumor (typically 3-fold lower) in combination with specific local genomic characteristics (e.g. lower average coverage due to for example local GC content or involving low complexity sequences) (Alioto et al., 2015).

While reconstruction of the derived chromosomal tumor genome topology based on the 68 truth set somatic SVs results in an overall stable genomic configuration for most derived chromatids harboring a single centromere and two telomeres, some breakpoint junctions are still clearly missing. This is corroborated by the fact that not for all copy number alterations breakpoint junctions were identified at either end. Our results indicate that these missing events typically involve centromeric regions that are not directly accessible by any current sequencing technology. Annotation data provided by the GRIDSS2 SV caller (Cameron et al., 2020). suggests a junction between a single break-end in chromosome 3 and the centromere in chromosome 1, which shows a copy number change. This can probably not be resolved directly due to the repeated nature of the centromeric region. When excluding the missing events that likely involve centromeres, there are 2 copy number aberrations that remain unexplained by the truth set, providing room for further improvement based on the existing or to be generated data.

Although this study was not designed to compare performance of sequencing platforms or data analysis pipelines, some interesting observations can be made. First, there is clear complementarity between the various platforms for the comprehensive identification of all real events. However, bioinformatic pipelines for somatic SV detection are still clearly in different stages for the different platforms with the most commonly used Illumina-based approaches yielding lowest numbers of false positives. We believe future tool optimisation for somatic SV calling, assisted by gold reference truth sets as well as the development of platform-specific germline and artefact filtering data sets (‘pools of normals’) based on large numbers of samples, will effectively address this challenge. Second, data analysis pipelines yield different annotations for the same event. This calls for further standardisation of variant annotation and nomenclature, although some observed differences are intrinsic to the use of short and long-read technologies. For example, a long templated insertion may be called as two independent translocations by short-read SV callers, while long read-based technology would detect this readily as an insertion. Third, despite previous studies showing the added value of long reads for SV detection for germline events, our somatic SV truth set is resolved almost in its entirety with the ILL short read dataset. While this may in part be due to the more advanced somatic SV calling pipelines developed for short-read data, this observation may also be explained by fundamental differences between germline and somatic SVs, where the latter are much more randomly distributed throughout the genome than inherited germline events. Germline variants more often involve complex or repetitive regions of the genome which might reflect mechanistic differences like for example the more frequent involvement of non-allelic homologous recombination, or be due to differences in selective pressure. As a consequence, somatic events may thus on average be more effectively detected.

The COLO829 cell line has the advantage that it is, in contrast to real tumor samples, a renewable source that can be used for assessing the impact of future platform developments or the performance of completely new technologies for somatic mutation detection. Although the COLO829 cell line is representative for structural variation as observed in cancer, including small and large copy number alterations (including aneuploidies) and both simple and complex SV events, it is not necessarily representative in all aspects for real tumor samples. First, tumor samples do typically not consist of tumor cells only but are a mix of tumor and normal cells (e.g. stromal cells and infiltrating immune cells). We show that the raw data obtained in this study can be used effectively to mimic variable tumor purity and that the truth set is instrumental for assessing the performance of the bioinformatic data analysis tools at variable tumor purity. As expected, our results show that both recall and precision heavily depend on tumor purity for all platforms. Secondly, tumors evolve continuously and are typically genetically heterogeneous, especially primary tumors, involving potentially subclonal SV events. While the COLO829 cell line has been shown to be genetically heterogeneous and evolving over time and thus could in principle capture this tumor feature properly, this variation is dynamic and might be variable between cell line isolates as already demonstrated by the various studies on this cell line (Craig et al., 2016; Velazquez-Villarreal et al., 2020) and thus limit the utility of a single defined truth set obtained as presented here. Finally, tumors are in general very heterogeneous both within the context of a specific tumor type, but especially between tumor types. For example, microsatellite instable (MSI) tumors show a high number of small indels (Fujimoto et al., 2020), homologous recombination deficient (HRD) tumors present many deletions with microhomology and large duplications (Nguyen et al., 2020) and paediatric haematological cancers cancers usually show very low mutational load but enhanced levels of somatic SVs, although often involving specific but complex genomic loci (e.g. the IgH locus) (Andersson et al., 2015; Ma et al., 2018). The specificity for capturing such heterogeneity effectively or the impact of specific genomic events that may co-occur in a given tumor sample, like for example whole genome duplication or chromothripsis, on overall performance of a specific sequencing technique or data analysis tool is of course not captured in a single cell line and requires the development of complementary datasets. Analysing additional cancer cell lines with matching normal cell lines provide an attractive route towards this goal as these represent in principle an endless source of genomic material for benchmarking of future DNA analysis technologies, but also for quality monitoring in routine production labs under accreditation. However, availability of suited cell lines that represent the full genetic diversity of cancer is a clear limitation. Ideally, one would thus resort to thoroughly analysed real tumor samples, even though in practice availability of sufficient material for multi-lab and multi-technology analyses can be problematic and sharing and reusing of patient material and data may require complex consenting and legal procedures.

Taken together, we believe the SV truth set described here as well as the underlying raw data, are a valuable resource for benchmarking and fine-tuning analysis settings of somatic SV calling tools, but the data may also be used for the development of novel analysis tools, for example phasing of structural variants. All analysis results and raw data are publicly available to enable such applications without access restrictions (ENA accession number: PRJEB27698 and an overview of the available data and specific access link can be found at **Supplementary Table 6).** We demonstrate this utility by analysing the impact of tumor purity and sequencing depth on SV recall and precision for different technologies, thereby providing valuable insights in the potential impact of technology platform choice and experimental design in relation to diagnostic accuracy and overall costs. Furthermore, these results highlight the need of benchmarking somatic SV detection methods at different tumor purities and sequencing depths rather than under a single fixed condition, since these parameters are highly variable within and between cohorts and can result in strong performance variation.

## Materials and Methods

### Sample source

COLO829 (ATCC^®^ CRL-1974™) and COLO829BL (ATCC^®^ CRL-1980™) cell lines were obtained from ATCC in September 2017. A single batch of cells was thawed and cells were expanded and grown according to standard procedures as recommended by ATCC. Cell pellets were split for technology-specific DNA isolation at 33 days (COLO829 & COLO829BL for the ILL and ONT datasets), 35 days (COLO829 for the PB, 10X and BNG datasets) and 23 days (COLO829BL for the PB, 10X and BNG datasets).

### Genomic analyses per technology

#### Illumina

COLO829 and COLO829BL libraries were prepped with Truseq Nano reagent kit and sequenced on the HiSeq X Ten platform using standard settings and reagent kits (chemistry version V2.5). Reads were mapped to GRCh37 with BWA mem (version 0.7.5, (Li, 2013)), followed by indel realignment with GATK (v3.4-46, (DePristo et al., 2011)). SVs were called jointly for COLO829 and COLO829BL with GRIDSS (v2.0.1,(Cameron et al., 2020)). Somatics SVs were filtered with the GRIDSS somatic SV filtering script (https://github.com/PapenfussLab/gridss/blob/master/scripts/gridss_somatic_filter.R).

#### Nanopore

COLO829 and COLO829BL libraries were sequenced on the MinION and GridION platforms using R9.4 flow cells. Reads were mapped to GRCh37 with NGMLR (v0.2.6, default parameters, (Sedlazeck et al., 2018)) with default parameters. SV calling was performed with both NanoSV (v. 1.2.2, default parameters, (Cretu Stancu et al., 2017)) and Sniffles (v1.0.9, parameters “--*report_BND --genotype*”, (Sedlazeck et al., 2018)) for COLO829 and COLO829BL separately. All SV calls for both NanoSV and Sniffles were merged with SURVIVOR (v1.0.6,(Jeffares et al., 2017)) with a distance of 200 bp and calls with evidence in COLO829BL for NanoSV or Sniffles were discarded.

#### PacBio

COLO829 and COLO829BL libraries were sequenced on the Sequel System with the 5.0 chemistry (binding kit 101-365-900; sequencing kit 101-309-500). Reads were mapped to GRCh37 with minimap2 (v2.11-r797, (Li, 2018)). SVs were called jointly for COLO829 and COLO829BL with pbsv (v2.0.1, https://github.com/pacificbiosciences/pbsv/) using default parameters. Somatic SV calls were filtered by removing any call with a supporting read in COLO829BL.

#### 10X

COLO829 and COLO829BL 10x genomics libraries were prepared on the Chromium platform and sequenced on the NovaSeq platform (chemistry version V1). Reads were analyzed with the LongRanger WGS pipeline (v2.2.2) separately for COLO829 (somatic mode) and COLO829BL (default parameters). SV calls for COLO829 and COLO829BL were merged with SURVIVOR (v. 1.0.6,(Jeffares et al., 2017)) with an overlap distance of 200 bp and SV calls with evidence in COLO829BL were discarded.

#### Bionano

DNA for COLO829 and COLO829BL was labelled using the Bionano Direct Label and Stain (DLS) kit. The labelled DNA was linearized in a Saphyr chip and imaging was performed on the Saphyr instrument. SV calling was performed on the Bionano Access platform. For each sample, 1.5 million cultured cells were used to purify ultra-high molecular weight DNA using the SP Blood & Cell Culture DNA Isolation Kit following manufacturer instructions (Bionano genomics, San Diego USA). Briefly, after counting, white blood cells were pelleted (2200g for 2mn) and treated with LBB lysis buffer and proteinase K to release genomic DNA (gDNA). After inactivation of proteinase K by PMSF treatment, genomic DNA was bound to a paramagnetic disk, washed and eluted in an appropriate buffer. Ultra-High molecular weight DNA was left to homogenize at room temperature overnight. The next day, DNA molecules were labeled using the DLS (Direct Label and Stain) DNA Labeling Kit (Bionano genomics, San Diego USA). Seven hundred and fifty nanograms of gDNA were labelled in presence of Direct Label Enzyme (DLE-1) and DL-green fluorophores. After clean-up of the excess of DL-Green fluorophores and rapid digestion of the remaining DLE-1 enzyme by proteinase K, DNA backbone was counterstained overnight before quantitation and visualization on a Saphyr instrument. A volume of 8.5 microliter of labelled gDNA solution of concentration between 4 and 12ng/ul was loaded on the Saphyr chip and scanned on the Saphyr instrument (Bionano genomics, San Diego USA). A total of 1.6 Tb and 1.5 Tb of data was collected for the cancer and blood sample, respectively.

De novo assembly Pipeline and Copy number variants calling were performed and against the Genome Reference Consortium Human Build 37 (GRCh37) HG19 human genome assembly (RefAligner version 7520). Events detected by the de novo assembly pipeline were subsequently compared against the matched blood control, and those that are absent in the assembly or the molecules of the control were considered as somatic variants (https://bionanogenomics.com/wp-content/uploads/2018/04/30190-Bionano-Solve-Theory-of-Operation-Variant-Annotation-Pipeline.pdf).

#### Consolidation of SV calls

Somatic SV calls for each dataset (ILL, ONT, PB and 10X) were merged using SURVIVOR (v. 1.0.6 (Jeffares et al., 2017) with an overlap distance of 200bp.

### Depth and molecular length calculations

Average base depth and depth distribution for ILL, ONT, PB and 10X was calculated based on 1,000,000 random positions on the genome with Sambamba (v0.6.5, (Tarasov et al., 2015)). Average base depth for BNG was calculated based on the same 1,000,000 random positions using Bedtools (v2.25.0, (Quinlan and Hall, 2010)).

Average molecular length and length distribution was calculated based on insert size for ILL, read length for ONT and PB, on synthetic molecular length based on the MI tag for 10X, on optical map length for BNG. For ILL, average insert size was calculated using Picard (v1.141, http://broadinstitute.github.io/picard).

### Copy number analysis

CNA calling was performed on the ILL dataset with BIC-SEQ2 (v0.7.2, (Xi et al., 2016)). For the remaining datasets, BAM and optical map (xmap) files were converted to BED format using Bedtools (v2.25.0, (Quinlan and Hall, 2010)) and CNA calling was performed with Ginkgo (Garvin et al., 2015). CNA calls from the different datasets were merged using 1MB bins to calculate Pearson’s correlation between datasets and for plotting.

### Validations

#### Capture

For each break-junction of the merged somatic SV calls 2 capture probes of 100 bp in length were designed, one at either side of the breakpoint, with a maximum distance of 100bp from the breakpoint at GC percentage as close as possible to 50%, for a total of 18148 custom probes. These custom capture probes were then ordered from Twist Biosciences. Then, libraries for COLO829 and COLO829BL were prepared and hybridized with the biotin-labelled custom targeted probes following the manufacturer’s protocol (Twist Biosciences catalog IDs: 100253, 100255, 100527, 100400). Using streptavidin beads the hybridized DNA was pulled from the DNA pool, and amplified by PCR. Enriched targeted libraries were sequenced on the Illumina NextSeq platform. NextSeq-Capture validation sequencing data were mapped with BWA mem (v0.7.5,(Li, 2013)) and SV calling was performed with Manta (v, REF), independently for COLO829 and COLO829BL. SV calls for COLO829 and COLO829BL were merged using SURVIVOR (v1.0.6, overlap distance of 50bp, (Jeffares et al., 2017)) and only calls with no evidence in COLO829BL were considered as somatic and validated.

#### PCR

We selected 88 high-confidence SV candidates for PCR validation based on an initial screening of the somatic SV truth set with IGV and added 296 randomly selected additional SV candidates for a total of 384 assays. We automatically designed primers for these SV breakpoints using Primer3 (v1.1.4, (Untergasser et al., 2012)). PCR assays were performed on COLO829 and COLO829BL genomic DNA. Libraries were prepared for PCR results and sequenced on both the MiSeq and ONT-MinION platforms. MiSeq-PCR validation sequencing data were mapped with BWA mem (v0.7.5,(Li, 2013)) and SV calling was performed with Manta (v0.29.5, (Chen et al., 2016)), independently for COLO829 and COLO829BL. ONT PCR validation sequencing data were mapped with minimap2 (v2.15, (Li, 2018)), and SV calling was performed with NanoSV (v1.2.2, default parameters, (Cretu Stancu et al., 2017)) independently for COLO829 and COLO829BL. Moreover, 70 additional SV calls that were shown as somatic in the Capture validation set were also subjected to PCR and products were sequenced on the MinION through the same protocol described above.

SV calls for COLO829 and COLO829BL from the Miseq-PCR and the two Nanopore-PCR sets were merged using SURVIVOR (v1.0.6, overlap distance of 50bp, (Jeffares et al., 2017)). Only SV calls with no evidence in any of the COLO829BL sets were considered somatic and validated.

#### FISH

For FISH validation, we selected probes that bind to 6 genomic regions, including Chromosome Enumeration Probes (CEP) for the centromeric region of chromosome 13, 16 and 18 (CEP13, CEP16, CEP18), labeled with SpectrumOrange (Abbott Vysis, Downers Grove, IL) and centromeric region of chromosome 9 (CEP9), labeled with SpectrumAqua (Leica Biosystems, Amsterdam). Furthermore, locus specific break-apart probes for chromosome 2p23 fusion (SpectrumOrange/SpectrumGreen, Vysis ALK Break Apart, Abbott Vysis, Downers Grove, IL) and 9p24 fusion (SpectrumOrange/SpectrumGreen Leica Biosystems, Amsterdam) were used.

COLO829 cells were dissociated using trypsin, counted, washed and diluted to contain a total of 50,000 cells in 100 μl. Monolayer cell suspensions were concentrated on a microscope slide using cytospin. Then, FISH was performed according to diagnostic standards. Slides were visualized on a Leica DM5500 fluorescence microscope and for each probe, 100 cells/slide were recorded.

### SV selection pipeline

Merged somatic SV calls were overlapped with the validation outcomes with SURVIVOR (v. 1.0.6, (Jeffares et al., 2017)) using an overlap distance of 50bp (PCR, CAPTURE) and 1kbp (BNG). Only somatic SV calls with support from multiple datasets and calls with support from a single dataset which were validated were selected. SVs involving unstable microsatellites were not considered as part of our analyses. All calls were manually curated by using the SV-plaudit cloud based framework (Belyeu et al., 2018) that uses Samplot to generate images from SV coordinates and BAM files. We generated such images for the somatic SV calls for each dataset (ILL, ONT, PB and 10x) and for the validations (PCR-ONT, PCR-MISEQ and CAPTURE). We evaluated each of these image datasets independently and classified each somatic SV call as “somatic”, “germline” or “false positive”. We also used the Integrated Genome Viewer (IGV, v2.4.0, (Robinson et al., 2017)) to verify some SVs. We performed the same analysis on 176 randomly selected SV calls belonging to a single dataset and which were not validated. Finally, we gathered the somatic SV calls and generated the final somatic VCF file.

### Comparison to external sources

CNA calls from (Arora et al., 2019) were downloaded (HiSeq dataset, https://www.nygenome.org/bioinformatics/3-cancer-cell-lines-on-2-sequencers/) and lifted to GRCh37 genomic coordinates with liftOver (UCSC). CNA calls from the four different single cell clusters were obtained from (Velazquez-Villarreal et al., 2020). These datasets were then merged using 1MB bins to calculate Pearson’s correlation between datasets and for plotting.

The two somatic SV sets from (Arora et al., 2019) (HiSeq and NovaSeq sets, https://www.nygenome.org/bioinformatics/3-cancer-cell-lines-on-2-sequencers/) were downloaded. Since these are BEDPE files based on GRCh38 genomic coordinates, we converted our somatic SV truth set to BEDPE format and lifted it to those coordinates using the liftOver tool from UCSC. We then intersected those SV sets with our truth set using Bedtools (v2.25.0, (Quinlan and Hall, 2010)) and differentiated between SVs with overlap on both sides, overlap only on one side and not overlapping. We lifted all SVs with no overlap or one-sided overlap and manually evaluated them in our data using IGV v2.4.0, (Robinson et al., 2017)).

### Tumor purity and sequencing depth analysis

For tumor purity simulations in each of the ILL, ONT and PB datasets, COLO829 and COLO829BL BAM files were randomly subsampled and mixed in different ratios, dependent on the sequencing depth to achieve *in silico* tumor purities of 10, 20, 25, 50 and 75 with Sambamba (v0.6.5, (Tarasov et al., 2015)). The same somatic SV calling pipeline used for the different datasets was applied to each of the tumor purity subsets. The resulting somatic SV file of each tumor purity subset was overlapped using a window of 100bp with the truth set VCF to determine the number of true and false positives and true negatives. This experiment was performed in triplicate for each tumor purity and each technology with the original COLO829 BAM file as positive control (100% tumor purity) and the original COLO829BL BAM file as negative control (0% tumor purity).

For sequencing depth simulations using the ILL dataset, one of the triplicates from each tumor purity simulation was selected together with the COLO829 and COLO829BL files. Each of these BAM files was subsampled to depths of 1x, 5x, 10x, 30x and 50x (plus the original 98x) with Sambamba (v0.6.5, (Tarasov et al., 2015)). Somatic SV calling was performed independently for each of the subsets and the resulting somatic SV VCF file was overlapped with the truth set to determine the number of true and false positives and false negatives.

## Data availability

Genomic data is available on EGA project PRJEB27698 Raw, somatic and truth set VCF files, and CNA files are available in Zenodo DOI: 10.5281/zenodo.3988185

## Code availability

All code used in the preparation of the somatic SV truth set is available at: https://github.com/UMCUGenetics/COLO829_somaticSV The code used for simulations of tumor purity and sequencing depth is available at: https://github.com/UMCUGenetics/tumps Figure panels 2-A, 3-C and 3-D were created using Biorender.com.

## Author contributions

*Conceptualization:* JEV-I, BY, RJAF, WPK, EC

*Provided material or generated data:* NJMB, EdB, MN, IR

*Data analysis:* JEV-I, DC, JE, JK, SvL, TM, PP, AWCP, MvR, AMW

*Validation:* NJMB, IR, MGMR, MvR

*Writing manuscript:* JEV-I, WPK, EC

*Project supervision:* WPK, EC

## Acknowledgements

We thank Pacific Biosciences and BioNano for their kind support generating and analysing data. JEV-I is supported by the Gieskes Strijbis Foundation (1816199). This work was performed as part of the EU-funded Horizon2020 EUCANcan project (funding to EC) and the Netherlands X-omics Initiative funded by NWO, project 184.034.019.

## Conflict of Interest Statement

AWCP is an employee of Bionano Genomics. AMW is an employee and shareholder of Pacific Biosciences.

## List of supplementary material

- **Supplementary Figures:**

○ Supplementary Figure 1: Copy number correlation within our datasets and external datasets.
○ Supplementary Figure 2: Generation of a somatic SV truth set
○ Supplementary Figure 3: Characterization of the somatic SV truth set
- **Supplementary Tables**

○ Supplementary Table 1: Dataset metrics
○ Supplementary Table 2: Manual curation results
○ Supplementary Table 3: Truth set annotated
○ Supplementary Table 4: CNAs annotated
○ Supplementary Table 5: Arora unique SVs annotated
○ Supplementary Table 6: Data accession details
- **Supplementary Movie**: Reconstruction of the breakage-fusion-bridge event in chromosome 3

**Supplementary Figure 1 - Related to figure 1:**
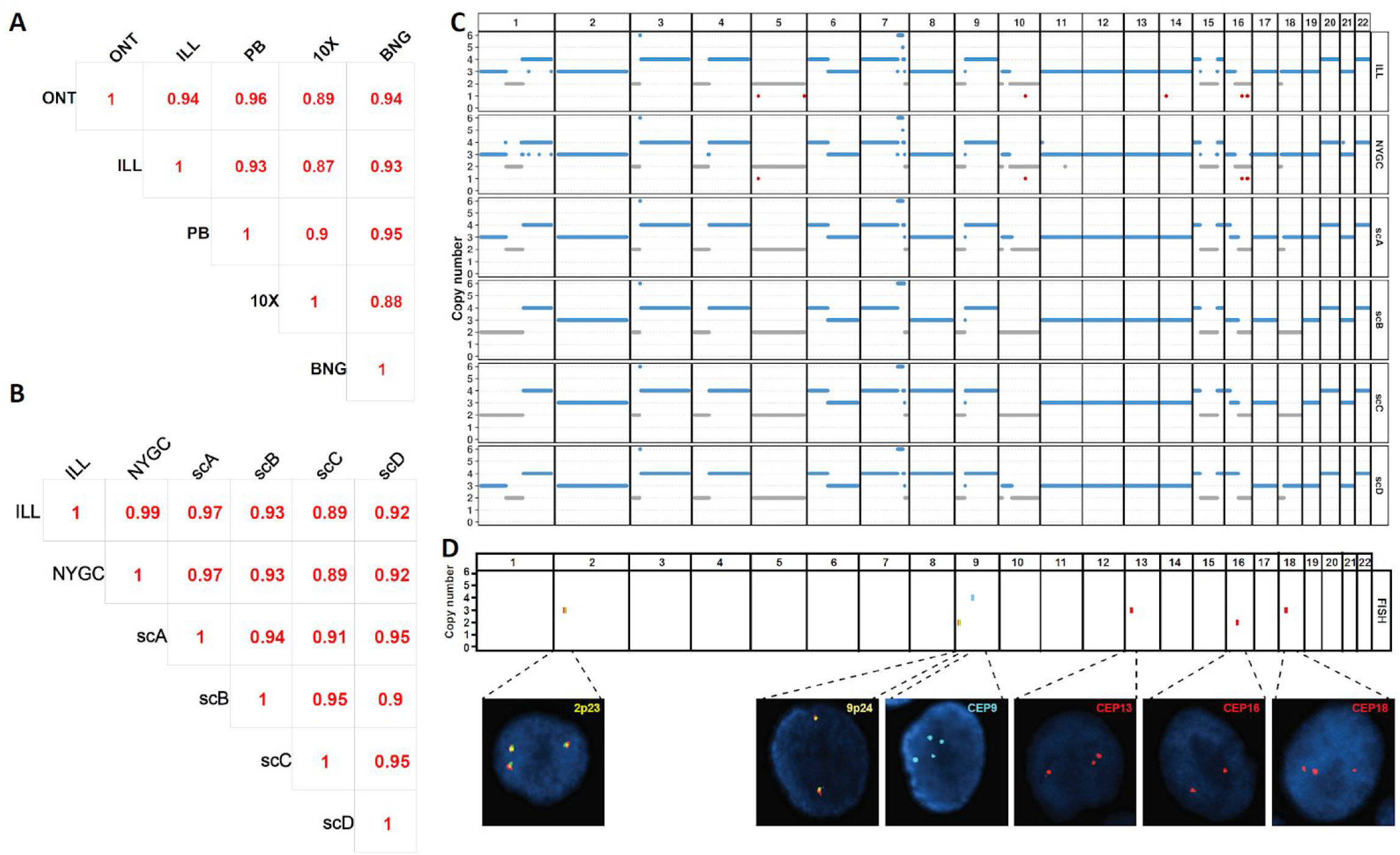
Copy number correlation within our datasets and external datasets. Correlation index of CNA calls for **(A)** each of the pairwise comparisons of the datasets generated in our study and **(B)** the comparison of our ILL dataset and the external sets from bulk sequencing in NYGC (Arora et al., 2019) and the 4 clusters differentiated by single cell sequencing (scA-D) (Arora et al., 2019; Velazquez-Villarreal et al., 2020). **(C)** Copy number profile of the ILL and the external sets. **(D)** Copy number status of 6 distinct genomic locations as determined by FISH

**Supplementary Figure 2 - Related to figure 2:**
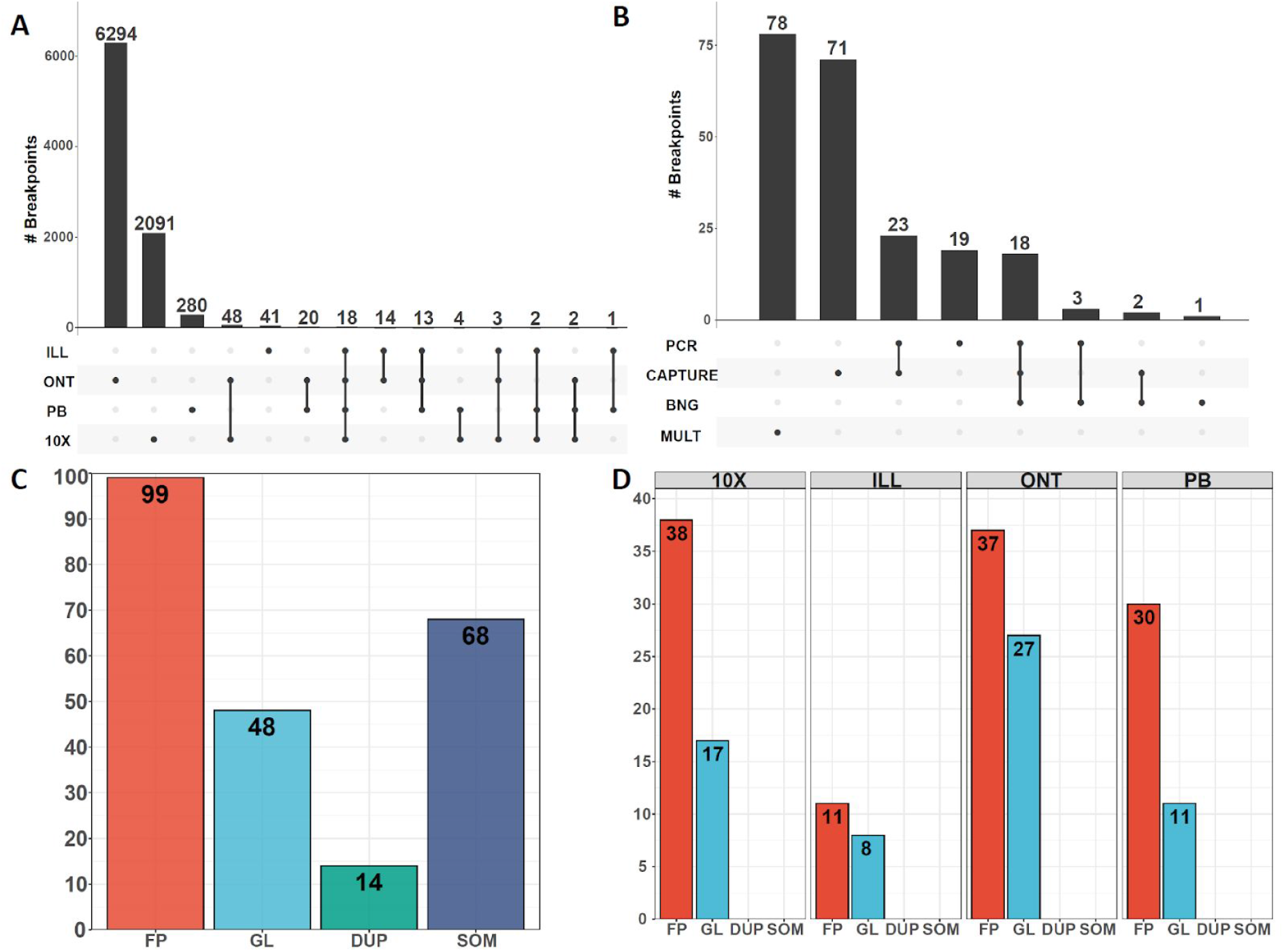
Generation of a somatic SV truth set. **(A)** Intersection of the total of 8,831 candidate SV calls merged from all platforms used and presence per in the raw call set per technology. **(B)** Number of validated somatic SV calls per validation approach including multi technology support (MULT). Manual curation statistics for **(C)** validated or multi-dataset SV calls and **(D)** non-validated and single-dataset SV calls. FP = false positive, GL = evidence in germline, DUP = duplication of an already called SV, SOM = real somatic variant.

**Supplementary Figure 3 - Related to figure 3:**
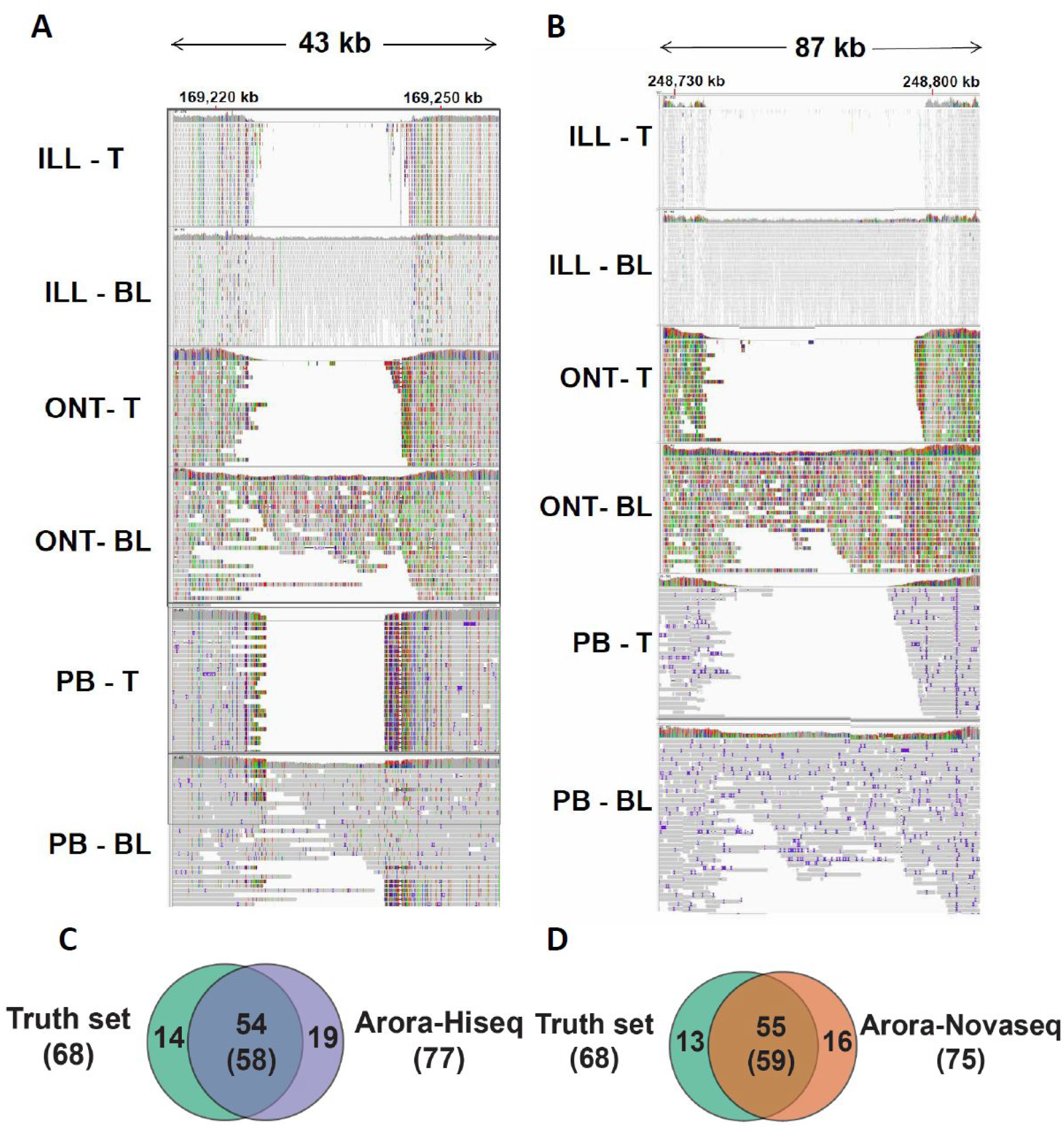
Characterization of the somatic SV truth set. **(A, B)** IGV screenshots of mapped reads from the ILL, ONT and PB datasets for COLO829 (T) and COLO829BL (BL) of two CNAs on chromosome 1 without associated somatic SVs in the truth set. Overlap of somatic SV calls between our truth set and the two somatic SV sets reported by (Arora et al., 2019), the Hiseq set **(C)** and the Novaseq set **(D)**. One-sided overlaps (i.e. when only one breakpoint of the SV overlaps) are included on the overlap. Numbers in parenthesis indicate the overlap from the Arora set point of view.

**Supplementary Movie - Reconstruction of the breakage-fusion-bridge event in chromosome 3:** Animated reconstruction of a breakage-fusion-bridge event consistent with the breakpoints and copy number profile of chromosome 3 in COLO829. This circos plot shows the evolution in time and over various cell divisions of the chromosome involving 4 cycles of breakage-fusion-bridge followed by a genome doubling event. The innermost track shows minor allele ploidy (orange indicates loss, blue amplification). The next track shows the copy number profile (purple indicates loss, green amplification). The line track shows the reconstructed chromosome. Breakpoints are represented by triangles and connecting arcs, telomeric ends of the chromosome by squares, and unrepaired double-stranded breaks by circles. DNA gained by replication and new breakpoints formed through DNA repair are indicated in blue, with lost DNA in orange. The outer track shows chromosome number and coordinate. A non-linear chromosomal coordinate scale is used with distances between breakpoints shown in black overlaying the copy number track. A cell cycle clock is shown in the upper left corner indicating at what point in the cell cycle each rearrangement occurs. The final stabilising repair to the centromere of another chromosome is omitted for clarity.

